# Epitopes targeted by T cells in convalescent COVID-19 patients

**DOI:** 10.1101/2020.08.26.267724

**Authors:** Ahmed A. Quadeer, Syed Faraz Ahmed, Matthew R. McKay

## Abstract

Knowledge of the epitopes of SARS-CoV-2 that are targeted by T cells in convalescent patients is important for understanding T cell immunity against COVID-19. This information can aid the design, development and assessment of COVID-19 vaccines, and inform novel diagnostic technologies. Here we provide a unified description and meta-analysis of emerging data of SARS-CoV-2 T cell epitopes compiled from 15 independent studies of cohorts of convalescent COVID-19 patients. Our analysis demonstrates the broad diversity of T cell epitopes that have been collectively recorded for SARS-CoV-2, while also identifying a selected set of immunoprevalent epitopes that induced consistent T cell responses in multiple cohorts and in a large fraction of tested patients. The landscape of SARS-CoV-2 T cell epitopes that we describe can help guide future immunological studies, including those related to vaccines and diagnostics. A web-based platform has been developed to help complement these efforts.

## Main text

Severe acute respiratory syndrome coronavirus 2 (SARS-CoV-2), the causative agent of COVID-19, has led to a global public health crisis. Development of COVID-19 vaccines and diagnostic tests are aided by an understanding of the natural protective immune responses against SARS-CoV-2. This includes both humoral and cellular immunity, mediated by antibodies and T cells respectively. Significant COVID-19 research has focused on understanding antibody responses (Wu et al., 2020; Yong et al., 2020; Zohar and Alter, 2020); though, studies informing the role of T cells have also started to emerge.

Initial results suggest the potential key role that T cells may play in protecting against COVID-19 (Grifoni et al., 2020a; Weiskopf et al., 2020). Studies of convalescent COVID-19 patients have detected SARS-CoV-2-specific T cells nine months post infection (Li et al., 2020), giving promising signs for the potential of T cells to provide lasting immunity. Longevity may be a concern for antibody responses, which have been reported to decline within just a few months post infection (Long et al., 2020; Seow et al., 2020; Snyder et al., 2020). Consistent observations have also been reported for the most closely related human coronavirus, SARS-CoV, for which T cells were shown to persist up to 17 years post infection (Le Bert et al., 2020), whereas antibody responses waned after a few years (Tang et al., 2011).

Characterizing SARS-CoV-2 T cell epitopes, as well as their human leukocyte antigen (HLA) association, is important for multiple reasons. It informs of the expected SARS-CoV-2 natural or vaccine-induced T cell responses in a population of specific ethnicity or a specific geographical region, which is tied to the composition of HLA alleles prevalent in that population. It can help in assessing the T cell responses expected to be induced by COVID-19 vaccines (which currently mainly focus on the spike (S) protein (Le et al., 2020)), while providing possible directions for boosting the T cell response by including specific immunodominant epitopes. It can guide vaccine assessment studies in probing whether a vaccine induces T cell responses similar to those commonly generated during natural infection. It can also aid in monitoring potential viral escape from T cell responses via genetic mutations, and can facilitate the development of T-cell-based diagnostics for distinguishing convalescent from unexposed individuals. T-cell-based diagnostics may have advantages over serological assays given the uncertainties related to the appearance and persistence of SARS-CoV-2-specific antibody responses in infected patients (Liu et al., 2020; Long et al., 2020; Seow et al., 2020; Snyder et al., 2020).

Here we present a unified account of the current knowledge (as of 20 December 2020) of SARS-CoV-2 T cell epitopes associated with convalescent COVID-19 patients. We collate and analyse data of T cell epitopes that have been identified experimentally in independent studies of blood samples from different patient cohorts. This data is compiled from 15 studies (ten published, five preprints) of T cell responses in a total of 848 convalescent COVID-19 patients (Table 1).

**Table 1:**
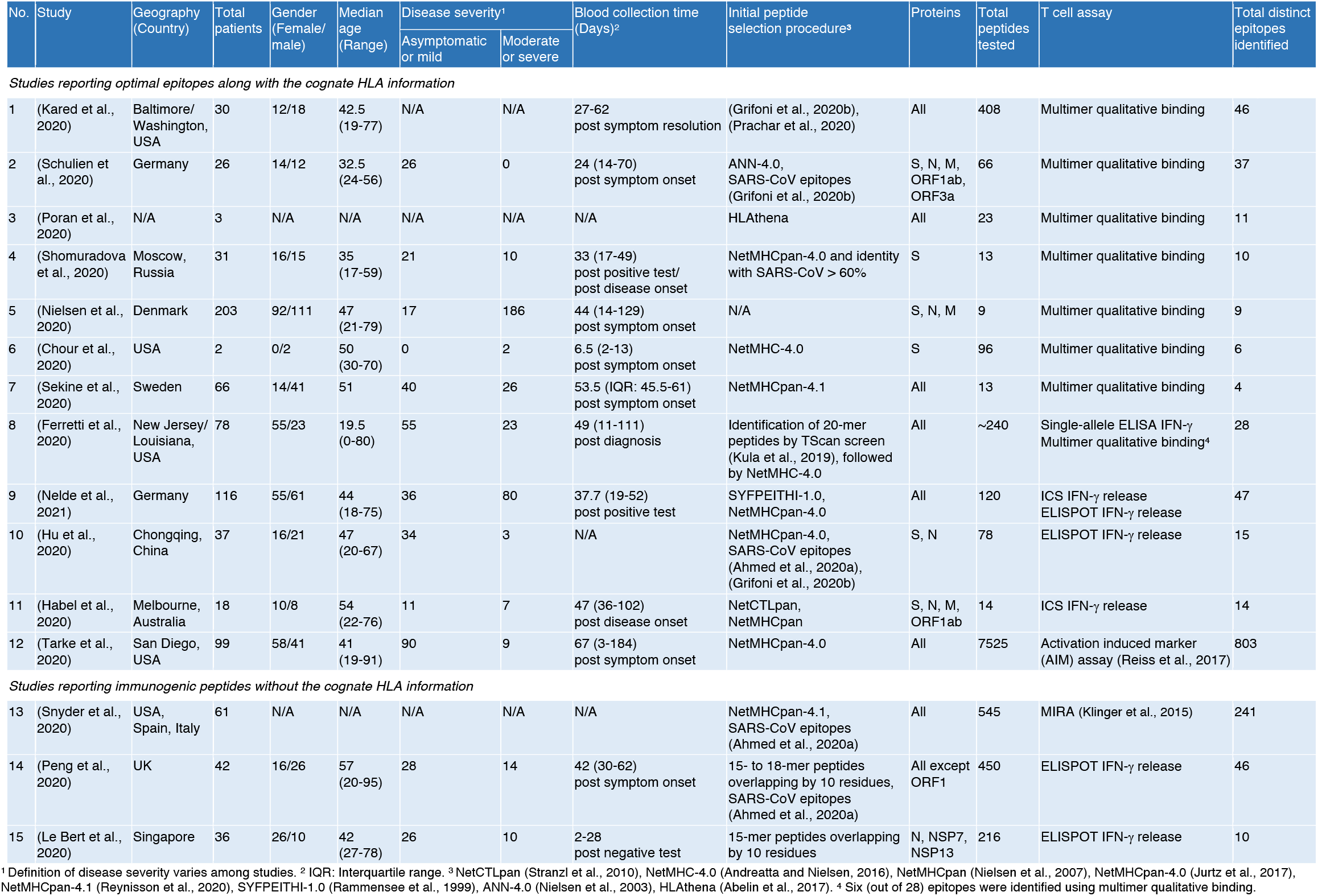
Summary of immunological studies reporting SARS-CoV-2 T cell epitopes targeted in convalescent COVID-19 patients (as of 20 December 2020).

| No. | Study | Geography<br>(Country) | Total<br>patients | Gender<br>(Female/<br>male) | Median<br>age<br>(Range) | Disease severity<sup>1</sup> |  | Blood collection time<br>(Days)<sup>2</sup> | Initial peptide<br>selection procedure<sup>3</sup> | Proteins | Total<br>peptides<br>tested | T cell assay | Total distinct<br>epitopes<br>identified |
| --- | --- | --- | --- | --- | --- | --- | --- | --- | --- | --- | --- | --- | --- |
|  |  |  |  |  |  | Asymptomatic<br>or mild | Moderate<br>or severe |  |  |  |  |  |  |
| Studies reporting optimal epitopes along with the cognate HLA information |  |  |  |  |  |  |  |  |  |  |  |  |  |
| 1 | (Kared et al., 2020) | Baltimore/<br>Washington,<br>USA | 30 | 12/18 | 42.5<br>(19-77) | N/A | N/A | 27-62<br>post symptom resolution | (Grifoni et al., 2020b),<br>(Prachar et al., 2020) | All | 408 | Multimer qualitative binding | 46 |
| 2 | (Schulien et al., 2020) | Germany | 26 | 14/12 | 32.5<br>(24-56) | 26 | 0 | 24 (14-70)<br>post symptom onset | ANN-4.0,<br>SARS-CoV epitopes<br>(Grifoni et al., 2020b) | S, N, M,<br>ORF1ab,<br>ORF3a | 66 | Multimer qualitative binding | 37 |
| 3 | (Poran et al., 2020) | N/A | 3 | N/A | N/A | N/A | N/A | N/A | HLAthena | All | 23 | Multimer qualitative binding | 11 |
| 4 | (Shomuradova et al., 2020) | Moscow,<br>Russia | 31 | 16/15 | 35<br>(17-59) | 21 | 10 | 33 (17-49)<br>post positive test/<br>post disease onset | NetMHCpan-4.0 and identity<br>with SARS-CoV > 60% | S | 13 | Multimer qualitative binding | 10 |
| 5 | (Nielsen et al., 2020) | Denmark | 203 | 92/111 | 47<br>(21-79) | 17 | 186 | 44 (14-129)<br>post symptom onset | N/A | S, N, M | 9 | Multimer qualitative binding | 9 |
| 6 | (Chour et al., 2020) | USA | 2 | 0/2 | 50<br>(30-70) | 0 | 2 | 6.5 (2-13)<br>post symptom onset | NetMHC-4.0 | S | 96 | Multimer qualitative binding | 6 |
| 7 | (Sekine et al., 2020) | Sweden | 66 | 14/41 | 51 | 40 | 26 | 53.5 (IQR: 45.5-61)<br>post symptom onset | NetMHCpan-4.1 | All | 13 | Multimer qualitative binding | 4 |
| 8 | (Ferretti et al., 2020) | New Jersey/<br>Louisiana,<br>USA | 78 | 55/23 | 19.5<br>(0-80) | 55 | 23 | 49 (11-111)<br>post diagnosis | Identification of 20-mer<br>peptides by TScan screen<br>(Kula et al., 2019), followed<br>by NetMHC-4.0 | All | ~240 | Single-allele ELISA IFN-γ<br>Multimer qualitative binding<sup>4</sup> | 28 |
| 9 | (Nelde et al., 2021) | Germany | 116 | 55/61 | 44<br>(18-75) | 36 | 80 | 37.7 (19-52)<br>post positive test | SYFPEITHI-1.0,<br>NetMHCpan-4.0 | All | 120 | ICS IFN-γ release<br>ELISPOT IFN-γ release | 47 |
| 10 | (Hu et al., 2020) | Chongqing,<br>China | 37 | 16/21 | 47<br>(20-67) | 34 | 3 | N/A | NetMHCpan-4.0,<br>SARS-CoV epitopes<br>(Ahmed et al., 2020a),<br>(Grifoni et al., 2020b) | S, N | 78 | ELISPOT IFN-γ release | 15 |
| 11 | (Habel et al., 2020) | Melbourne,<br>Australia | 18 | 10/8 | 54<br>(22-76) | 11 | 7 | 47 (36-102)<br>post disease onset | NetCTLpan,<br>NetMHCpan | S, N, M,<br>ORF1ab | 14 | ICS IFN-γ release | 14 |
| 12 | (Tarke et al., 2020) | San Diego,<br>USA | 99 | 58/41 | 41<br>(19-91) | 90 | 9 | 67 (3-184)<br>post symptom onset | NetMHCpan-4.0 | All | 7525 | Activation induced marker<br>(AIM) assay (Reiss et al., 2017) | 803 |
| *Studies reporting immunogenic peptides without the cognate HLA information* |  |  |  |  |  |  |  |  |  |  |  |  |  |
| 13 | (Snyder et al., 2020) | USA, Spain, Italy | 61 | N/A | N/A | N/A | N/A | N/A | NetMHCpan-4.1, SARS-CoV epitopes (Ahmed et al., 2020a) | All | 545 | MIRA (Klinger et al., 2015) | 241 |
| 14 | (Peng et al., 2020) | UK | 42 | 16/26 | 57 (20-95) | 28 | 14 | 42 (30-62) post symptom onset | 15- to 18-mer peptides overlapping by 10 residues, SARS-CoV epitopes (Ahmed et al., 2020a) | All except ORF1 | 450 | ELISPOT IFN-γ release | 46 |
| 15 | (Le Bert et al., 2020) | Singapore | 36 | 26/10 | 42 (27-78) | 26 | 10 | 2-28 post negative test | 15-mer peptides overlapping by 10 residues | N, NSP7, NSP13 | 216 | ELISPOT IFN-γ release | 10 |

<sup>1</sup> Definition of disease severity varies among studies. <sup>2</sup> IQR: Interquartile range. <sup>3</sup> NetCTLpan (Stranzl et al., 2010), NetMHC-4.0 (Andreatta and Nielsen, 2016), NetMHCpan (Nielsen et al., 2007), NetMHCpan-4.0 (Jurtz et al., 2017), NetMHCpan-4.1 (Reynisson et al., 2020), SYFPEITHI-1.0 (Rammensee et al., 1999), ANN-4.0 (Nielsen et al., 2003), HLAthena (Abelin et al., 2017). <sup>4</sup> Six (out of 28) epitopes were identified using multimer qualitative binding.

The set of convalescent patients collectively cover a population across four continents and are well distributed in age, gender, disease severity, and blood collection time, within and across studies (Table 1). Half of the studies have characterized T cell responses against the whole proteome, while the others have focused on responses mounted against subsets of SARS-CoV-2 proteins, typically involving the S protein. In the majority of cases, the T cell response was measured in blood samples of patients against a set of peptides predicted by bioinformatic tools or earlier bioinformatic-based analyses (Ahmed et al., 2020a; Grifoni et al., 2020b; Prachar et al., 2020), while a few studies employed overlapping peptide pools spanning the SARS-CoV-2 proteome. The majority of studies (no. 1-12 in Table 1) reported optimal epitopes along with cognate HLA information. Of these, seven studies determined HLA restrictions of the reported epitopes experimentally using multimer qualitative binding. Of the remaining five studies, one study (no. 8 in Table 1) employed mono-allelic cell line assays to identify specific HLA-restricted epitopes, while others (no. 9-12 in Table 1) inferred HLA restrictions using standard functional assays (such as activation induced marker (AIM) or ICS/ELISPOT IFN-γ release) and HLA haplotype information of patients. While the latter set of studies involved some predictive element in deconvolving from the patient haplotype the HLA allele responsible for the observed response, the reported HLA associations were supported by either testing HLA binding assays or ruling out the possibility of presentation by other alleles in the tested patients using accurate peptide-HLA binding prediction methods. The few studies that did not provide HLA information (no. 13-15 in Table 1) essentially reported responses against synthetic peptide pool libraries using functional assays, or molecular assays such as MIRA, and identified long immunogenic peptides. As complete information of epitopes was not available in these studies, we focus henceforth on the data reported by studies that provide optimal epitopes along with their HLA restriction (epitopes reported in studies 1-12 and two optimal epitopes reported along with their cognate HLA in study no. 14).

A total of 620 unique T cell epitopes with the information of cognate HLA allele have been reported in the immunological studies (see Methods for details). Of these, 544 are CD8^+^ (HLA class I restricted) and 76 are CD4+ (HLA class II restricted) (Figure 1A). The set of epitopes cover each protein of SARS-CoV-2, except ORF7b and ORF10. The largest number of epitopes fall within S and ORF1a—the most exposed protein and the longest SARS-CoV-2 ORF respectively. These epitopes broadly cover almost the entire S protein including the receptor-binding domain, while the majority of epitopes in ORF1a lie within the nsp3 (PL2-PROPapain-like proteinase) and nsp4 proteins (Supplementary Figure S1). These two proteins together participate in the assembly of virally-induced cytoplasmic double-membrane vesicles that are necessary for viral replication (UniProt, 2020). All identified epitopes have high genetic conservation (>0.9) among the ~200,000 SARS-CoV-2 sequences (as of 20 December 2020), except for seven epitopes (Figure 1B). Of these, three epitopes (_612_YQ<u>D</u>VNCTEV_620_ in S, _4706_NVLFSTVFP<u>P</u>TSFGP_4720_ and _4710_STVFP<u>P</u>TSF_4718_, in ORF1b) have a very low conservation (~0.09), since they encompass mutations (S:D614G and ORF1b:P4715L; underlined) that define a globally prevalent SARS-CoV-2 clade (Nextstrain (Hadfield et al., 2018), https://nextstrain.org). The genetic variation in three other epitopes (_211_NLVRDLPQGFS<u>A</u>LEP_225_ and _216_LPQGFS<u>A</u>L_223_ in S and _219_L<u>A</u>LLLLDRL_227_ in N; conservation 0.83) is attributed to mutations (S:A222V and N:A222P) that belong to a fast growing subclade in Europe, while variation in the last epitope (_57_<u>Q</u>SASKIITL_65_ in ORF3a; conservation 0.77) is attributed to the Q57H mutation that belongs to multiple SARS-CoV-2 subclades prevalent in North America and Europe. Of the identified mutations, S:D614G has been confirmed to increase viral fitness (Plante et al., 2020), while potential evolutionary advantages of the other mutations remain unclear. Our data, in general, provides little evidence to suggest that the observed genetic variation in SARS-CoV-2 may be attributed to escape from T cell immunity, though escape may become an important evolutionary factor once strong and diverse selective pressures are imposed by widespread vaccination, and this requires close monitoring.

**Figure 1.**
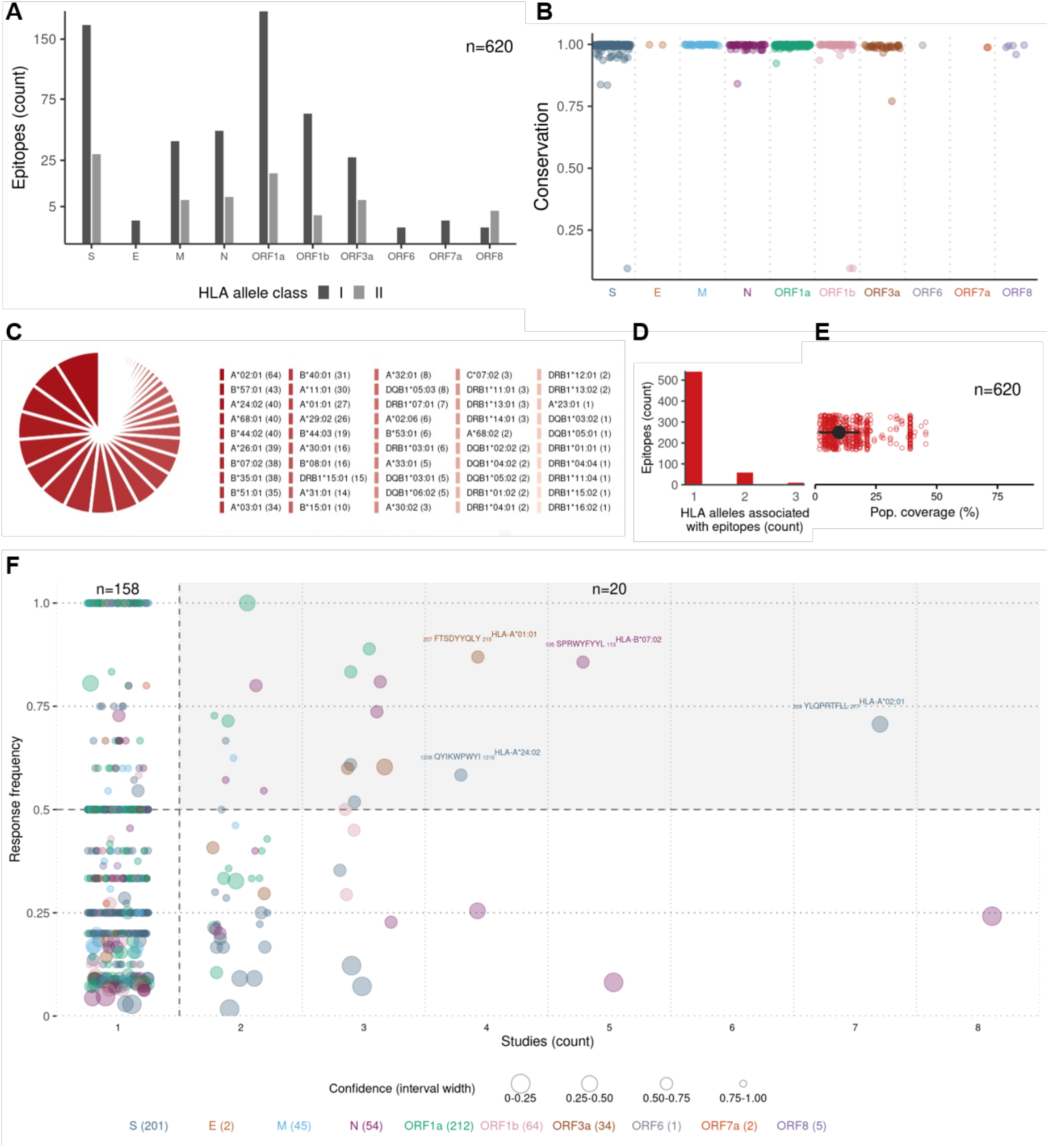
Features of SARS-CoV-2-specific T cell epitopes reported to elicit immune response in blood samples of convalescent COVID-19 patients, and identification of immunoprevalent epitopes. (**A**) The number of epitopes (n=620) according to HLA class restriction. (**B**) The conservation of each epitope among the global SARS-CoV-2 genetic sequences (as of 20 December 2020). (**C**) Diversity of HLA associations reported for SARS-CoV-2 epitopes. The number of epitopes associated with a particular HLA allele is shown in brackets. (**D**) The number of HLA alleles associated with each epitope. (**E**) Estimate of the global population coverage of each epitope (Methods). The median is shown as black circle. (**F**) Response frequency of unique epitope-HLA pairs versus the number of immunological studies reporting T cell response against them. The size of each circle represents the confidence in the respective RF value (Methods). The 20 immunoprevalent epitope-HLA pairs (having RF > 0.5 and reported in more than one study) are shown with shaded background, while the four highly immunoprevalent epitopes (having RF > 0.5 and reported in at least four studies) are labelled.

The reported epitopes have been associated to a total of 50 HLA class I and II alleles (Figure 1C). Most HLA alleles are associated with multiple epitopes, with 20 alleles each having an association with 10 epitopes or more. The same epitope may be presented by multiple HLA alleles, as evidenced by studies of the related SARS-CoV (Ahmed et al., 2020a) as well as other viruses (Vita et al., 2019). However, for the majority of SARS-CoV-2 epitopes, only a single associated HLA allele has been reported so far (Figure 1D). This appears due, in part, to the limited number of convalescent COVID-19 patients that have been studied (Table 1). The limited number of associated HLA alleles translates to a median global population coverage estimate per epitope of only 10% (Figure 1E). Thus, investigation of additional HLA alleles associated with the identified SARS-CoV-2 epitopes is required for providing a more accurate indication of their individual population coverage. An expanded list of likely HLA associations may be predicted for some of the reported epitopes by using prior knowledge of genetically-matched experimentally-determined T cell epitopes of SARS-CoV and their associated HLA alleles (for details, see Supplementary Table S1 and Supplementary Figure S2). These predictions, once confirmed, would provide an increase in the median population coverage of the selected SARS-CoV-2 epitopes from 16.8% to 40%, with a few epitopes having around 60% coverage (Supplementary Figure S2). Such predicted epitope-HLA pairs are recommended as targets for investigation in further immunological studies.

Quantifying the immunodominance of each reported epitope-HLA pair using the standard response frequency (RF) metric (Dhanda et al., 2018; Tian et al., 2019; Weiskopf et al., 2013) (Methods) revealed that 178 of the reported (710) epitope-HLA pairs had an RF score exceeding 0.5, indicating that they induced T cell responses in over half of the subjects tested across studies (Figure 1F). Confidence in the estimated RF values varies with the number of tested subjects, with higher confidence being attributed to epitope-HLA pairs with larger numbers of tested subjects (Methods). The majority (158/178) of epitope-HLA pairs with RF>0.5 had been reported in a single study only. Among these, pairs with high confidence appear promising, though responses against them should be investigated in different cohorts to further confirm their immunodominance. Responses against the remaining (20/178) epitope-HLA pairs were registered in more than one study, and while per-study variation was observed in the results (Supplementary Figure S3), these pairs appear to be immunoprevalent (Sidney et al., 2020). This is because responses against each epitope-HLA pair were recorded in more than half of the tested convalescent patients collectively across multiple studies, despite differences in characteristics of donor cohorts (age, gender, geographical location, disease severity), blood collection time, and methodology used for determining the epitopes (initial peptide selection procedure, T cell assay) (Table 1).

Interestingly, among the set of identified immunoprevalent epitope-HLA pairs (n=20), 35% had an identical match to experimentally-determined SARS-CoV epitope-HLA pairs. This fraction of matched epitope-HLA pairs is over three times higher than that (11.6%) in the complete set of SARS-COV-2 epitope-HLA pairs (P<10^-3^, Fisher’s exact test), and suggests that many of the immunoprevalent T cell epitopes of SARS-CoV-2 are also cross-reactive to SARS-CoV. Of the 20 identified immunoprevalent epitopes-HLA pairs, six belonged to N, five to S, five to ORF1a, three to ORF3a and one to M. This indicates that ~75% of the epitopes lie in proteins other than S, suggesting that vaccine candidates targeting these proteins may have benefits in terms of T cell immunity. Of the identified immunoprevalent epitopes, four epitopes (HLA-A*02:01-restricted _269_YLQPRTFLL_277_ in S, HLA-B*07:02-restricted _105_SPRWYFYYL_113_ in N, HLA-A*01:01-restricted _207_FTSDYYQLY_215_ in ORF3a, and HLA-A*24:02-restricted _1208_QYIKWPWYI_1216_ in S) appeared highly immunoprevalent, eliciting T cell responses in more than ~60% of the tested convalescent COVID-19 patients in four immunological studies or more. Collectively, around 71% of the global population is estimated to carry the associated HLA alleles, and hence may generate a T cell response against at least one of these four epitopes.

The SARS-CoV-2 T cell epitope data that we compile and report here has been integrated into a web-based dashboard (Ahmed et al., 2020b). This dashboard provides exportable data tables listing the SARS-CoV-2 epitopes, and graphical displays to summarize different characteristics of the epitopes, including aggregate information as well as specific details of the individual epitopes. We plan to update the dashboard with new experimental information as it becomes available, with the goal of aiding further research in understanding T cell responses against SARS-CoV-2, and in guiding studies related to COVID-19 vaccines and diagnostics. While we focused the current study on SARS-CoV-2-specific T cell responses recorded in convalescent patients, knowledge of T cell epitopes targeted in animal studies (Corbett et al., 2020; Smith et al., 2020) is also informative. These may inform potential immune targets in COVID-19-infected individuals, and can help in guiding further immunological experiments that seek to probe T cell responses that arise due to natural infection, or those elicited by vaccination. The SARS-CoV-2 epitopes reported to be targeted in animal models have also been incorporated into the web dashboard.

Overall, the data that we have described based on recent experimental studies demonstrates an impressive and diverse list of SARS-CoV-2 T cell epitopes targeted by convalescent COVID-19 patients. Subsets of these epitopes exhibit desirable properties, including high genetic conservation and high response frequency across multiple cohorts, and they appear to have the potential to collectively induce a T cell response in a large fraction of the population. Current knowledge of the landscape of T cell epitopes for SARS-CoV-2 is still evolving however, and further studies of different cohorts of convalescent patients, encompassing a broad diversity of HLA profiles, are required to provide a more comprehensive understanding. Moreover, further systematic studies are required to ascertain possible correlates between the responses against T cell epitopes and disease protection. The knowledge of SARS-CoV-2 T cell epitopes could play an important role in contributing to the fight against COVID-19 by guiding diverse applications and novel technologies, including the development, assessment and monitoring of vaccines, and the development of improved diagnostic assays.

## Acknowledgements

The authors were supported by the General Research Fund of the Hong Kong Research Grants Council (RGC) [Grant No. 16204519 and 16201620]. S.F.A. was additionally supported by the Hong Kong Ph.D. Fellowship Scheme (HKPFS).

## Author contributions

A.A.Q. and M.R.M. conceptualized the study and wrote the manuscript. A.A.Q. compiled the data from the literature. A.A.Q., S.F.A., and M.R.M. analyzed the data. A.A.Q. and S.F.A performed the computations. S.F.A. generated the figures and developed the web-dashboard.

## STAR METHODS

### KEY RESOURCES TABLE

| REAGENT or RESOURCE | SOURCE | IDENTIFIER |
| --- | --- | --- |
| **Deposited Data** |  |  |
| Epitopes targeted by T cells in convalescent COVID-19 patients: (Chour et al., 2020) | (Chour et al., 2020) | Figure 3 |
| Epitopes targeted by T cells in convalescent COVID-19 patients: (Shomuradova et al., 2020) | (Shomuradova et al., 2020) | Table 1 |
| Epitopes targeted by T cells in convalescent COVID-19 patients: (Nelde et al., 2021) | (Nelde et al., 2021) | Extended Data Tables 2 and 3 |
| Epitopes targeted by T cells in convalescent COVID-19 patients: (Poran et al., 2020) | (Poran et al., 2020) | Figure 3 |
| Epitopes targeted by T cells in convalescent COVID-19 patients: (Ferretti et al., 2020) | (Ferretti et al., 2020) | Table 1 |
| Epitopes targeted by T cells in convalescent COVID-19 patients: (Le Bert et al., 2020) | (Le Bert et al., 2020) | Table 1 |
| Epitopes targeted by T cells in convalescent COVID-19 patients: (Peng et al., 2020) | (Peng et al., 2020) | Tables 1 and 2 |
| Epitopes targeted by T cells in convalescent COVID-19 patients: (Snyder et al., 2020) | Data deposited by authors of (Snyder et al., 2020) in the Immune Epitope Database (IEDB; https://www.iedb.org/) | REF ID: 1036945 |
| Epitopes targeted by T cells in convalescent COVID-19 patients: (Kared et al., 2020) | (Kared et al., 2020) | Figure S2 |
| Epitopes targeted by T cells in convalescent COVID-19 patients: (Schulien et al., 2020) | (Schulien et al., 2020) | Table S1 |
| Epitopes targeted by T cells in convalescent COVID-19 patients: (Habel et al., 2020) | (Habel et al., 2020) | Figure 2B |
| Epitopes targeted by T cells in convalescent COVID-19 patients: (Hu et al., 2020) | (Hu et al., 2020) | Table 1 |
| Epitopes targeted by T cells in convalescent COVID-19 patients: (Nielsen et al., 2020) | (Nielsen et al., 2020) | Figure 5B |
| Epitopes targeted by T cells in convalescent COVID-19 patients: (Tarke et al., 2020) | (Tarke et al., 2020) | Tables S3 and S5 |
| Epitopes targeted by T cells in convalescent COVID-19 patients: (Sekine et al., 2020) | (Sekine et al., 2020) | Table S2 |
| Response frequencies of SARS-CoV-2 T cell epitopes | This paper | Table S1 |
| Genetic conservation of SARS-CoV-2 T cell epitopes among SARS-CoV-2 sequences (as of 20 December 2020) | This paper | Table S1 |
| Total subjects tested for each SARS-CoV-2 T cell epitope across studies | This paper | Table S1 |
| Total subjects responded to each SARS-CoV-2 T cell epitope across studies | This paper | Table S1 |
| Additional HLA alleles predicted for SARS-CoV-2 T cell epitopes | This paper | Table S1, Figure S2 |
| SARS-CoV-2 genome sequence data for conservation analysis | GISAID (https://www.gisaid.org) | All full genome and high coverage sequences available as of 20 December 2020 |
| SARS-CoV-2 reference genome for aligning the sequences | GenBank (https://www.ncbi.nlm.nih.gov/nuccore/NC_045512.2) | ID: NC_045512.2 |
| SARS-CoV T cell epitope data | IEDB; https://www.iedb.org/ | IEDB was queried for epitopes with positive MHC binding and positive T cell assays using "Severe acute respiratory syndrome-related coronavirus" as "Organism" on 21 February 2020. |
| **Software and Algorithms** |  |  |
| SARS-CoV-2 T cell epitope web-dashboard | This paper | https://www.mckayspcb.com/SARS2TcellEpitopes/ |
| MAFFT | (Katoh and Standley, 2013) | https://mafft.cbrc.jp/alignment/software/ |
| Estimated population coverage of SARS-CoV-2 T cell epitopes based on associated HLA alleles | IEDB Analysis Resource - Population coverage tool | http://tools.iedb.org/population/download/ |

### RESOURCE AVAILABILITY

#### Lead Contact

Further information and requests for resources should be directed to and will be fulfilled by the Lead Contact Matthew R. McKay.

#### Materials Availability

This study did not generate new materials.

#### Data and Code Availability

Compiled data of the SARS-CoV-2 T cell epitopes is available to download from the web dashboard (Ahmed et al., 2020b).

## METHOD DETAILS

### Epitope data

All information related to the epitopes was obtained from the literature or IEDB as reported in the Key Resource Table. A total of 1,327 epitopes were obtained from 15 immunological studies (Table 1). Removing the epitopes with no HLA allele information and those for which the number of tested and responded patients was not reported at a distinct epitope-HLA level resulted in a total of 620 unique epitopes (710 unique epitope-HLA pairs; listed in Supplementary Table S1). These corresponded to epitopes reported in studies 1-12 and two epitopes from study no. 14 (Table 1).

### Sequence data, epitope conservation and coverage

SARS-CoV-2 genomic sequences were obtained from the GISAID database (https://www.gisaid.org/) on 20 December 2020. We downloaded only the complete (full-genome) sequences derived from human hosts with high coverage using the options provided on the GISAID database. All of the 201,542 downloaded sequences were aligned to the SARS-CoV-2 reference genome (GenBank ID: NC_045512.2) using MAFFT (Katoh and Standley, 2013). The genomic MSA was translated using an in-house code to obtain the protein MSAs. The positions of the open reading frames provided with the reference sequence were used to identify the respective protein regions of the full genome.

The conservation of each SARS-CoV-2 T cell epitope was calculated as the fraction of SARS-CoV-2 sequences that encompassed the precise epitope sequence. The coverage of SARS-CoV-2 T cell epitopes at any position of a protein was calculated by counting the number of epitopes that included that position.

### Response frequency (RF)

RF score (Dhanda et al., 2018) was used to quantify the immunodominance of the SARS-CoV-2 epitopes reported to be recognized by T cells in convalescent COVID-19 patients. The RF score of an epitope is defined as follows:

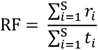

where *r_i_* is the number of subjects responding to the epitope in study *i, t_i_* is the number of subjects tested for a response against the epitope in study *i*, and *S* is the total number of studies. An RF score calculated using a large number of tested subjects would be more reliable than one calculated using relatively few subjects. To account for this, we computed the 95% confidence interval for the RF score of each epitope using the binomial cumulative distribution function (Dhanda et al., 2018). In Figure 1F, we defined the confidence in RF value of an epitope as the inverse of the length of the corresponding 95% confidence interval. That is, values of RF with a smaller 95% confidence interval have higher confidence, and vice versa.

### Estimating global population coverage of epitopes

The global population coverage of an epitope refers to the percentage of individuals in the world population that is expected to mount a T cell response against that epitope. The population coverage of a T cell epitope was calculated based on the HLA alleles associated with it using the tool downloaded from the IEDB Analysis Resource (http://tools.iedb.org/population/download/). This tool employs global HLA allele frequency data obtained from the Allele Frequency Net Database (http://www.allelefrequencies.net/) to estimate the population coverage.

## Supplemental Information

### Supplemental Tables

**Table S1** (included as a separate file). **Complete list of unique SARS-CoV-2-specific T cell epitope-HLA pairs (n=710) reported to elicit response in blood samples of convalescent COVID-19 patients.** It includes, for each epitope, the associated HLA allele, location within the proteome, genetic conservation in global SARS-CoV-2 sequences (as of 20 December 2020), and the computed response frequency across studies (as reported in Figure 1F). Number of total subjects tested, number of responding subjects, and response frequency per study for each epitope-HLA pair are also listed. For (n=79) SARS-CoV-2 epitopes having a genetic match with SARS-CoV known epitopes, the IEDB IDs of the corresponding SARS-CoV epitopes as well as the associated HLAs are listed. This list of epitopes is compiled from both published studies and non-peer-reviewed studies available so far only on preprint servers. Results from non-peer-reviewed studies should be interpreted with caution.

### Supplemental Figures

**Figure S1.**
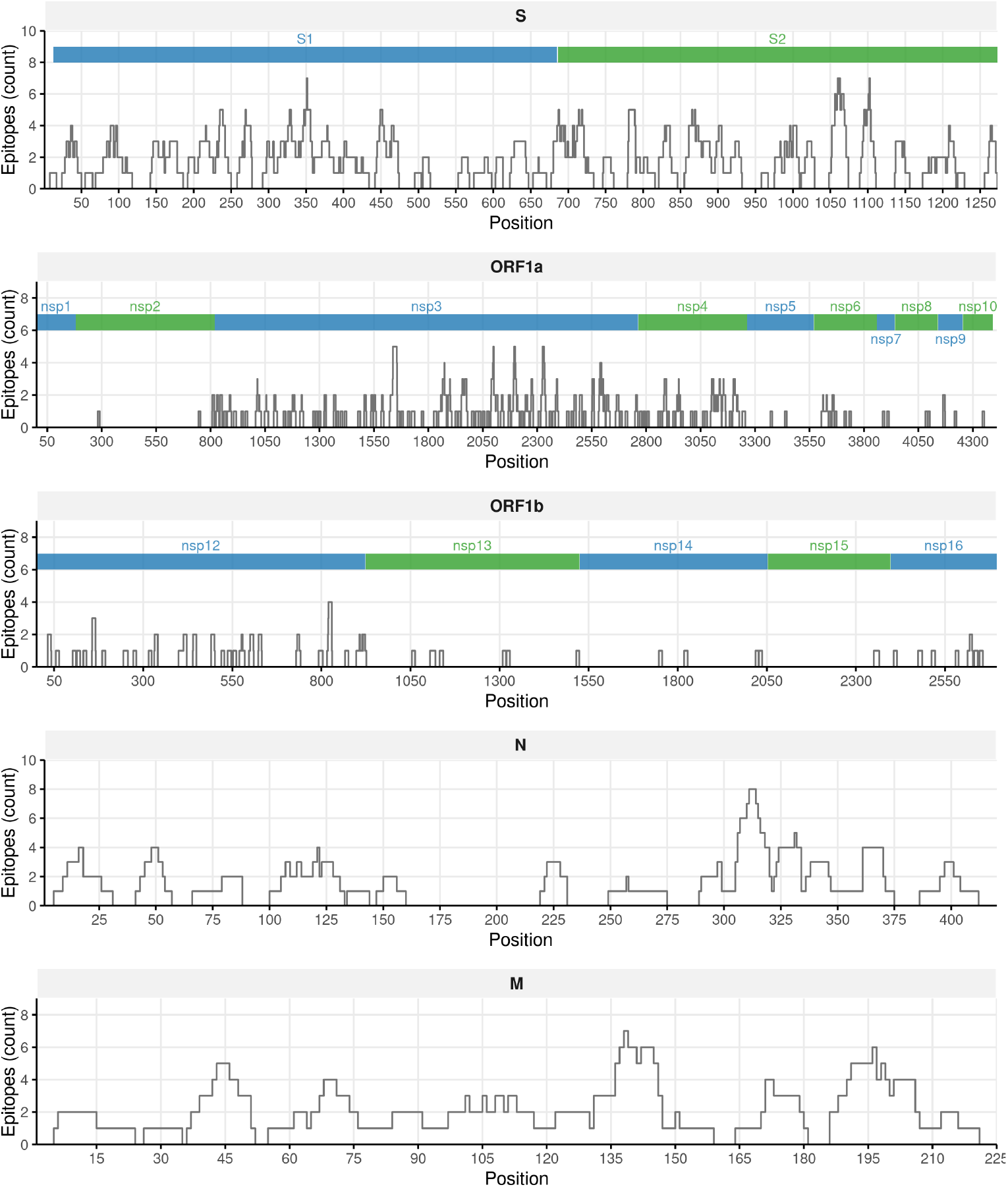
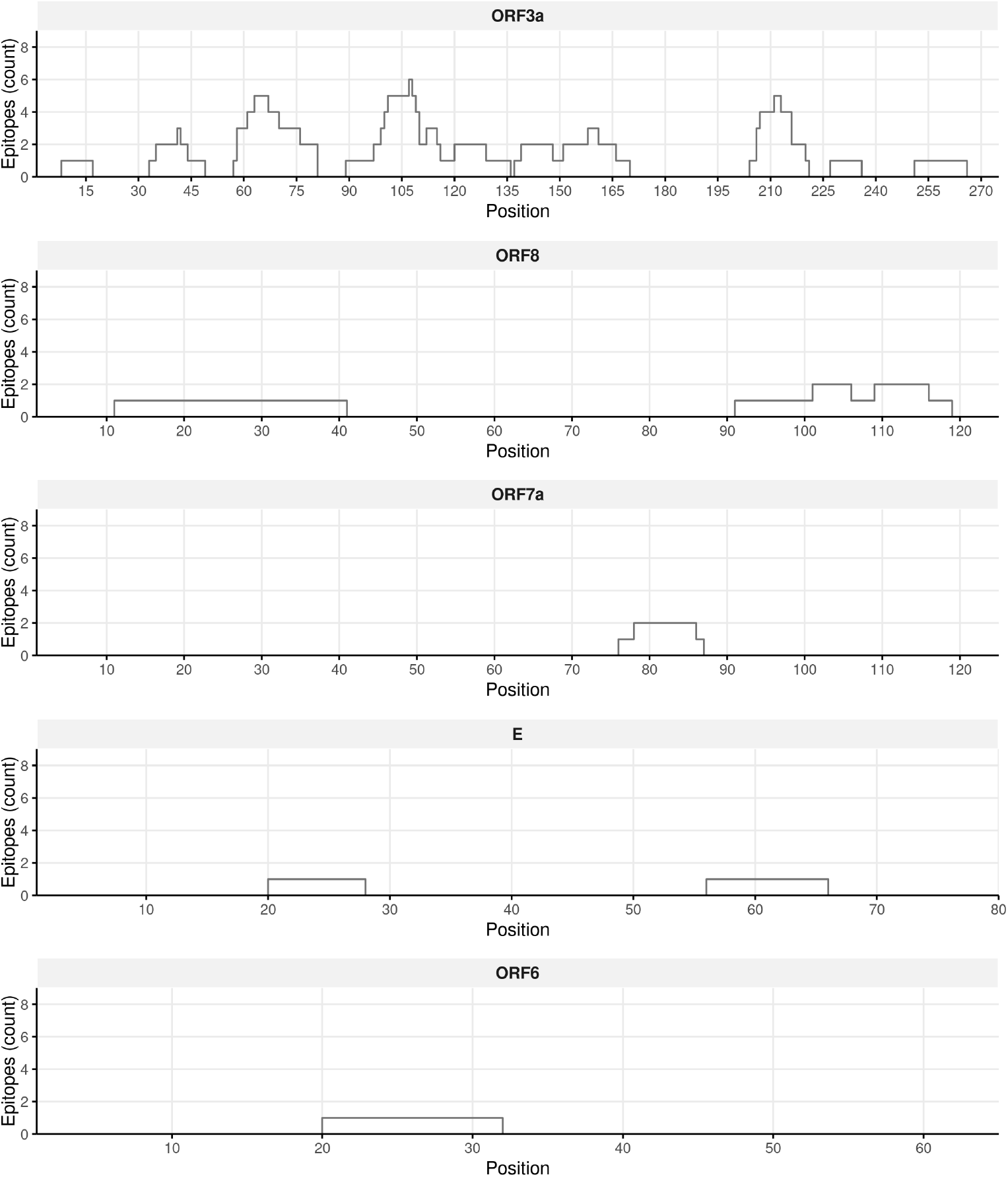
Epitope coverage of the SARS-CoV-2 proteins. Number of epitopes that cover the protein are indicated along its primary structure.

**Figure S2.**
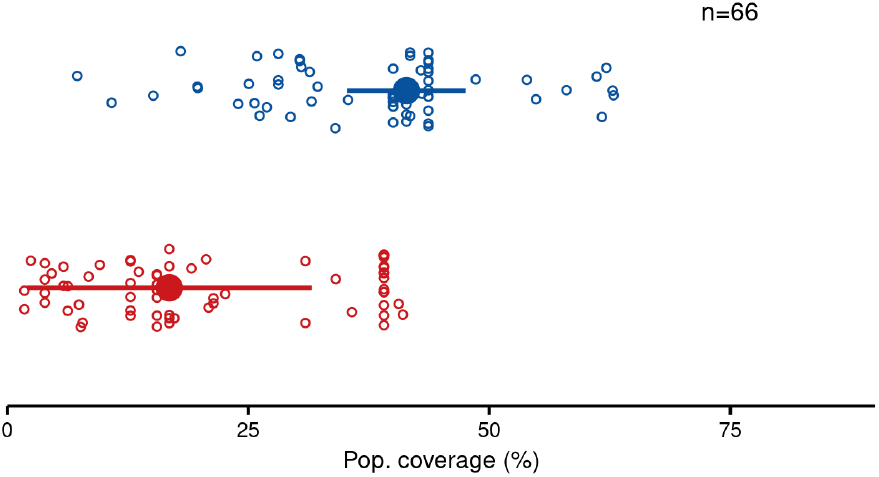
Estimated global population coverage of SARS-CoV-2 epitopes after augmenting HLA alleles associated with genetically-matched SARS-CoV epitopes. For 79 of the 620 experimentally-determined SARS-CoV-2 T cell epitopes, both epitope sequence and associated HLA alleles matched exactly with those of experimentally-determined SARS-CoV epitopes. This suggests that an epitope-HLA pair associated with a T cell response for one virus may very likely be associated with a response for the other (Košmrlj et al., 2008; Ngono and Shresta, 2019), and hence any additional HLA associations known for SARS-CoV epitopes (Ahmed et al., 2020c, 2020a) could augment the limited information available for SARS-CoV-2. Based on this rationale, additional HLA alleles from SARS-CoV data were identified for 66 of the 79 genetically-matched SARS-CoV-2 epitopes. For these 66 epitopes, the median population coverage significantly increased from 16.8% to 41.4% after augmenting HLA alleles from SARS-CoV data. Specific SARS-CoV-2 epitopes, such as _66_FPRGQGVPI_74_ in N and _171_ATSRTLSYY_179_ in M, were estimated to cover a high percentage of global population (~60%) after data augmentation (Table S1).

**Figure S3.**
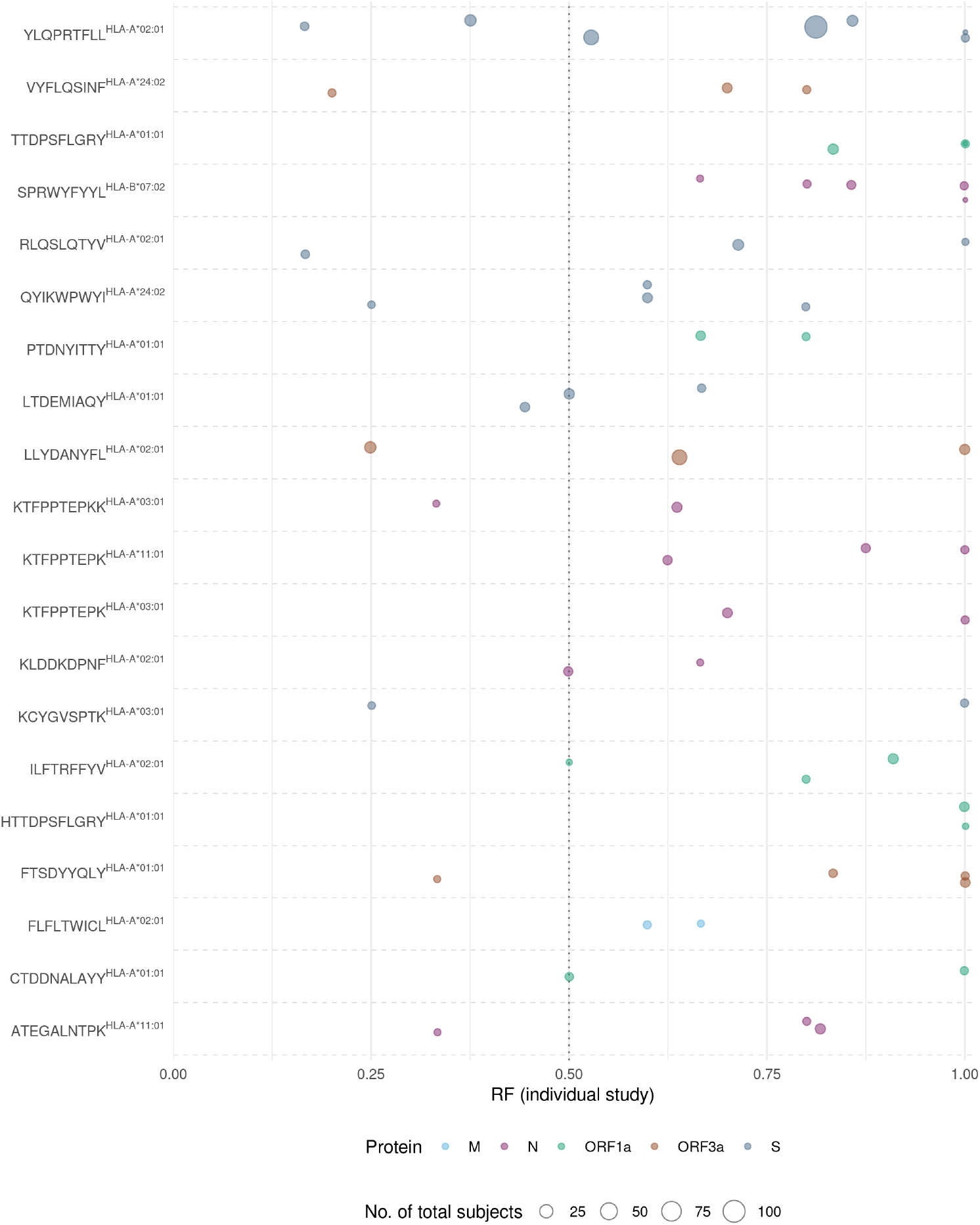
Response frequencies (RF) of immunoprevalent epitope-HLA pairs per individual study. For the 20 immunoprevalent epitope-HLA pairs (having RF > 0.5 and reported in more than one study) (Figure 1F), the RF computed per study is represented by circles. The size of these points indicates the total number of subjects that were tested for the specific epitope-HLA pair in a study. Note that each epitope-HLA pair in this set has RF > 0.5 in most of the immunological studies that reported a response against it.

